# Deep learning models of RNA base-pairing structures make accurate zero-shot predictions of base-base interactions of RNA complexes

**DOI:** 10.1101/2023.09.26.559463

**Authors:** Mei Lang, Thomas Litfin, Ke Chen, Jian Zhan, Yaoqi Zhou

## Abstract

The intricate network of RNA-RNA interactions, crucial for orchestrating essential cellular processes like transcriptional and translational regulation, has been unveiling through high-throughput techniques and computational predictions. With the emergence of deep learning methodologies, the question arises: how do these cutting-edge techniques for base-pairing prediction compare to traditional free-energy-based approaches, particularly when applied to the challenging domain of interaction prediction via chain concatenation? In this study, we employ base pairs derived from three-dimensional RNA complex structures as the gold standard benchmark to assess the performance of 23 different methods, including recently developed deep learning models. Our results demonstrate that the deep-learning-based methods, SPOT-RNA can be generalized to make accurate zero-shot predictions of RNA-RNA interactions not only between previously unseen RNA structures but also between RNAs without monomeric structures. The finding underscores the potential of deep learning as a robust tool for advancing our understanding of these complex molecular interactions.

## Introduction

Recent advancements in high-throughput techniques have unveiled a complex network of RNA-RNA interactions (RRIs) critical for governing transcriptional and translational processes. These interactions are pivotal in the biogenesis of various RNA molecules, including mRNAs, rRNA, tRNA, microRNAs, and circRNAs^1–3^. Large-scale detection of RRIs has been achieved through innovative approaches that combine cross-linking techniques with high-throughput sequencing. Notably, techniques such as PARIS^4^, SPLASH^5^, LIGR-seq^6^, and COMRADES^7^ have employed psoralen or its derivatives, as well as formaldehyde in the case of RIC-Seq^8^, for cross-linking. While these methods hold promise, they are not without limitations stemming from probe biases and ligation efficiencies ^1–3^. For example, KARR-seq, PARIS, and RIC-seq can only achieve about 55% true positive rate at 20% false positive rate for human 18S rRNA^5^. Furthermore, many of these high-throughput techniques have yet to achieve the single nucleotide resolution.

Attaining the nucleotide-level resolution in RNA structures has historically relied on traditional structure-determination methods such as X-ray crystallography, Nuclear Magnetic Resonance (NMR), and Cryo-electron microscopy. Yet, compared to proteins, determining RNA structures presents formidable challenges due to the unique physiochemical properties of nucleotides and the inherent fragility of RNA structures^9^. This is reflected from the fact that only a meagre 3% of structures in the Protein Data Bank contain RNAs, with even fewer dedicated to RNA-RNA complexes (681 as of March 16, 2023, before redundancy removal)^10^. This stark contrast becomes even more pronounced when considering the extensive collection of more than 31 million noncoding RNA sequences catalogued in the RNAcentral database ^11^. Given the cost and challenges associated with experimental approaches, there is an imperative need for development of complementary computational prediction techniques.

The existing methods for predicting RNA-RNA interactions (RRIs) can be broadly classified into alignment-based, free-energy-based, and homology modeling approaches^12,13^. Alignment-based techniques, such as GUUGle^14^ and RIsearch^15^, focus on inter-RNA base pairs while overlooking potential intra-RNA interactions. Free-energy-based methods can be categorized into those considering only intermolecular interactions for expediency (such as RNAhybrid^16^, RNAduplex^17^, RNAplex-c^18^, and DuplexFold^19^), those factoring in intramolecular interactions based on solvent accessibility (such as RNAup^20^, IntaRNA^21^, RNAplex-a^22^, and AccessFold^23^), and those accommodating both intra- and inter-molecular base pairs through sequence concatenation (such as PAIRFOLD^24^, RNAcofold^25^ and biFold^19^) or without restrictions (such as RactIP^26^). Homology-based techniques, exemplified by TargetRNA2^27^, CopraRNA^28^, RNAaliduplex^17^ and PETcofold^29^, utilize evolutionary information to infer binding.

Presently, ‘de novo’ RRI prediction methods predominantly rely on free-energy-based approaches, limited by their approximate energy or scoring functions, akin to the challenges faced in RNA secondary structure prediction^30^. Recent advancements have seen the emergence of deep learning-based methods, starting with SPOT-RNA^31^, which achieved the first end-to-end prediction of intra-RNA base pairs. Subsequent developments include mxfold2^32^, UFold^33^, and 2dRNA^34^. To further enhance prediction accuracy, SPOT-RNA2^35^ was developed to integrate evolutionary profiles and mutational coupling data generated by RNAcmap^36^.

In this study, we conducted a comprehensive benchmark of various methods for predicting inter-RNA interactions. Our evaluation encompassed traditional energy-based techniques and newly developed deep learning models based on simple sequence concatenation. To ensure a rigorous assessment, we employed base pairs derived from experimentally-determined RNA-RNA complex structures and eliminated monomer structures employed for training of SPOT-RNA and SPOT-RNA2 through a strict structural similarity cutoff (TM-score^37^ <0.3), in addition to sequence similarity cut off by CD-HIT. This challenging set of the complex of unseen structures revealed significant improvements of SPOT-RNA’s performance over the other 22 methods evaluated, underscoring the transferability of deep learning from intra-RNA to inter-RNA interaction prediction. The performance by SPOT-RNA is robust against different types of RNA complexes. Further tests on training-family-excluded test sets confirm the robustness of SPOT-RNA in heterodimeric complexes of RNAs, in particular.

## Results

### Method Comparison on Inter-RNA Base Pair Prediction

In this study, we compared 23 different RRI predictors on a benchmark set of 64 RNA-RNA pairs after excluding all monomer structures remotely similar to the RNA structures employed in the training and validation sets for SPOT-RNA and SPOT-RNA2. We evaluated their performance in untrained inter-RNA base pair prediction through precision/recall curves and F1 score distributions, as shown in Figure 1. The performance metrics, including overall F1-score, Mathews’ correlation coefficient (MCC) values, and the median F1-score and standard deviation of individual RNA pairs, are also summarized in Table 1. Predictors with probabilistic outputs are represented by precision-recall (PR) curves, while others are represented as single points. For all methods with chain concatenation for predicting RNA-RNA interactions, a low case “c” is appended to the method name. They are RNAfoldc, UFoldc, MXfold2c, EternaFoldc, SPOT-RNAc, and SPOT-RNA2c. No linker was employed because adding a 3-nucleotide link did not lead to performance improvement (See Methods).

**Figure 1:**
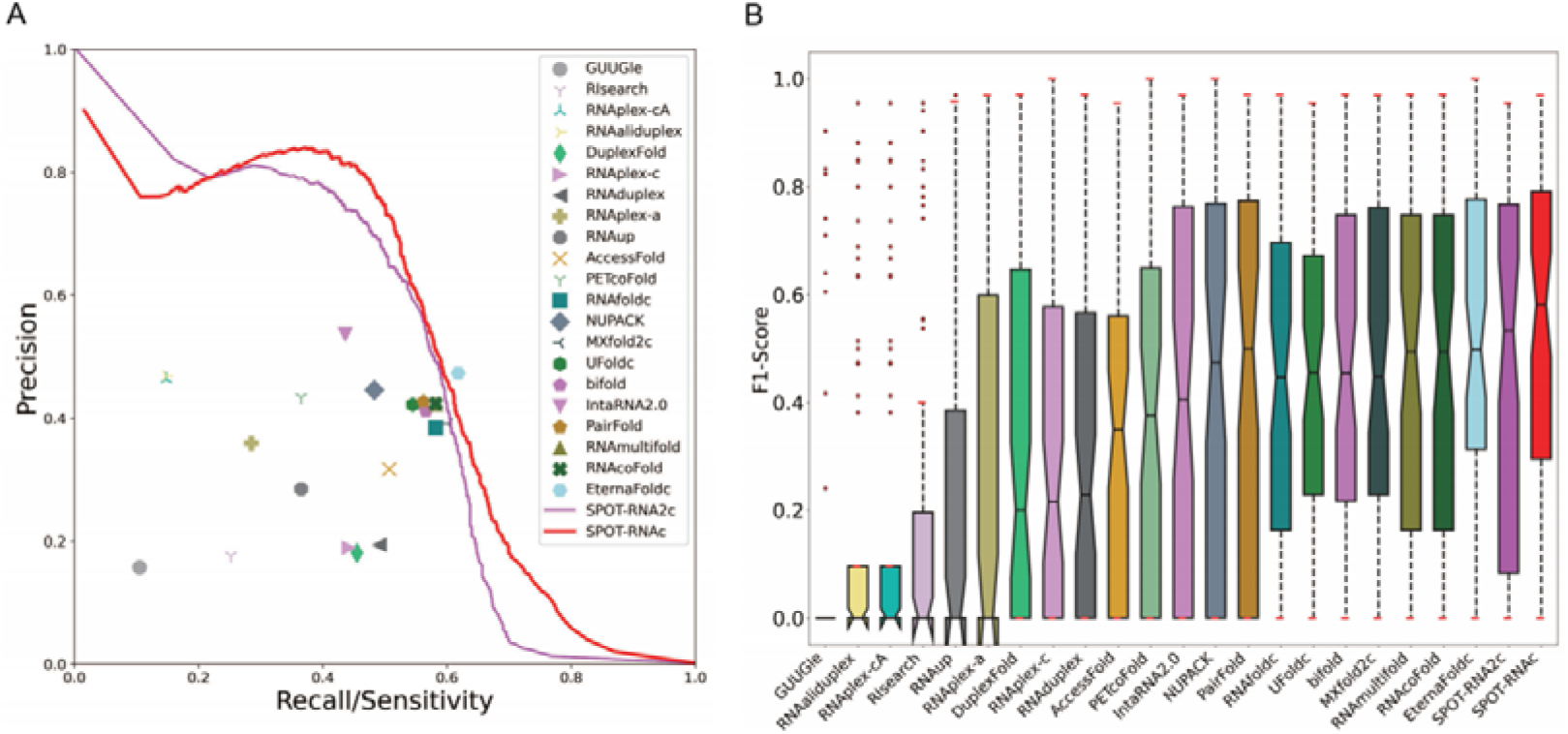
Performance comparison of 23 methods for inter-RNA base-pair prediction on the 64 complexes of RNA structures unseen by SPOT-RNA and SPOT-RNA2. All structures in the test set have the structural similarity score TM-Score<0.3 compared to the monomeric structures used in training and validating SPOT-RNA and SPOT-RNA2 methods. (A) Precision-recall curves (for those methods with probabilistic outputs) or points given by 23 methods (B) Distribution of F1 scores for inter-RNA base pair prediction for individual RNA pairs by the same 23 methods. Each boxplot shows the median, 25th, and 75th percentiles, with outliers represented by “•”. SPOT-RNAc exhibits the best performance for both overall and individual measures of F1-scores.

**Table 1:**
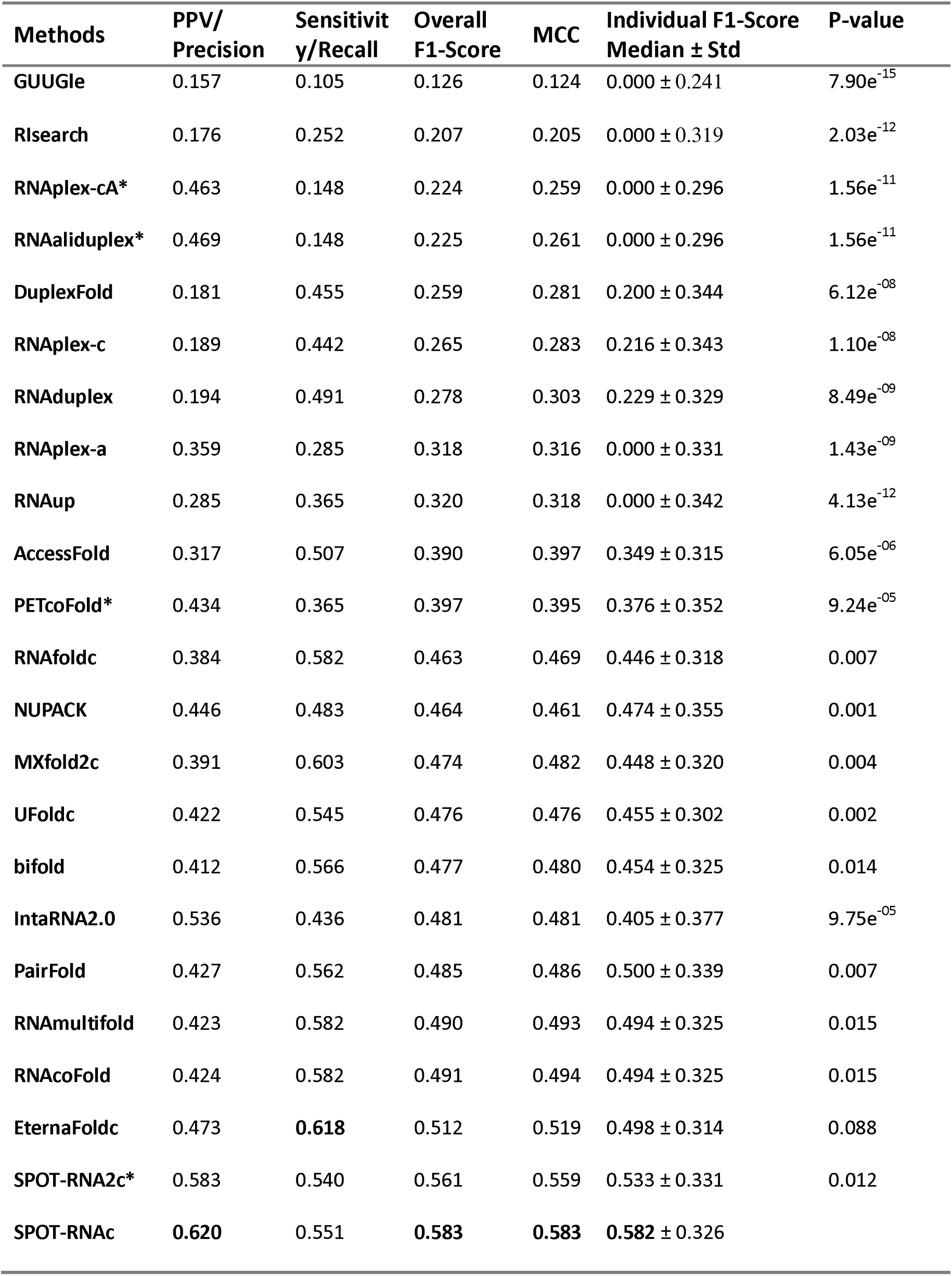
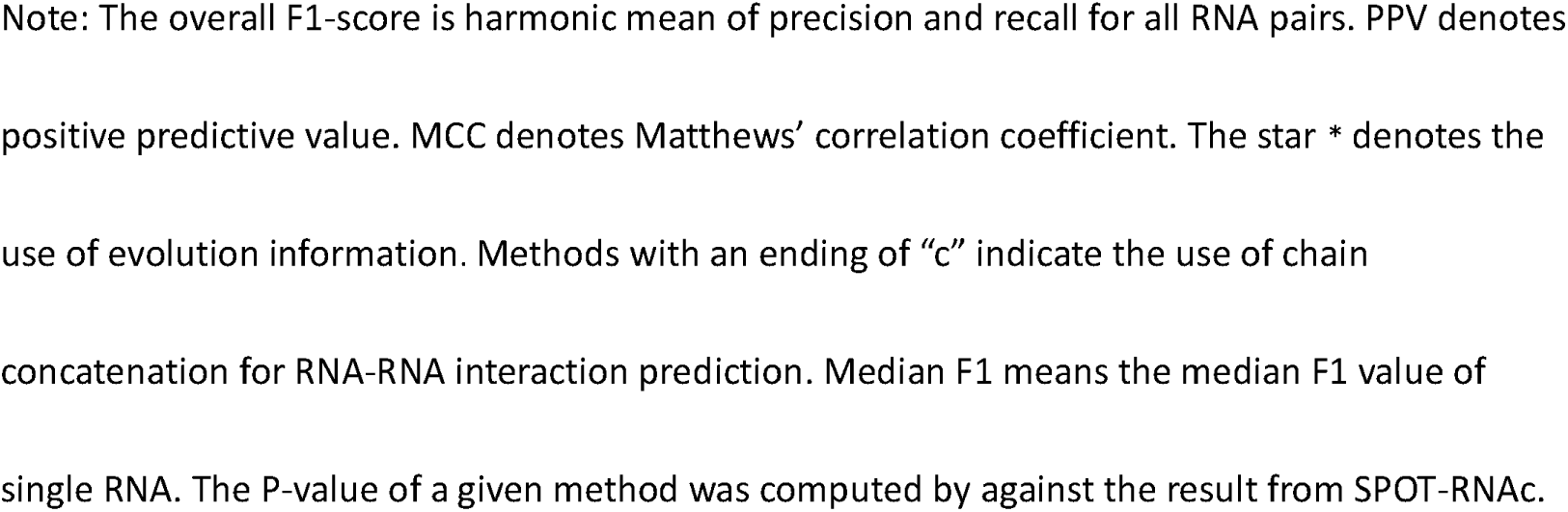
Performance Comparison of 23 Predictors of Inter-RNA Base Pairs on 64 Complexes of RNA Structures Unseen by SPOT-RNA and SPOT-RNA2. All single-chain structures in the test set have the structural similarity score TM-Score<0.3 compared to the monomeric structures used in training and validating SPOT-RNA and SPOT-RNA2 methods.

| Methods | PPV/<br>Precision | Sensitivity/<br>Recall | Overall<br>F1-Score | MCC | Individual F1-Score<br>Median ± Std | P-value |
| --- | --- | --- | --- | --- | --- | --- |
| GUUGle | 0.157 | 0.105 | 0.126 | 0.124 | 0.000 ± 0.241 | 7.90e<sup>-15</sup> |
| Rlsearch | 0.176 | 0.252 | 0.207 | 0.205 | 0.000 ± 0.319 | 2.03e<sup>-12</sup> |
| RNAplex-cA* | 0.463 | 0.148 | 0.224 | 0.259 | 0.000 ± 0.296 | 1.56e<sup>-11</sup> |
| RNAaliduplex* | 0.469 | 0.148 | 0.225 | 0.261 | 0.000 ± 0.296 | 1.56e<sup>-11</sup> |
| DuplexFold | 0.181 | 0.455 | 0.259 | 0.281 | 0.200 ± 0.344 | 6.12e<sup>-08</sup> |
| RNAplex-c | 0.189 | 0.442 | 0.265 | 0.283 | 0.216 ± 0.343 | 1.10e<sup>-08</sup> |
| RNAduplex | 0.194 | 0.491 | 0.278 | 0.303 | 0.229 ± 0.329 | 8.49e<sup>-09</sup> |
| RNAplex-a | 0.359 | 0.285 | 0.318 | 0.316 | 0.000 ± 0.331 | 1.43e<sup>-09</sup> |
| RNAup | 0.285 | 0.365 | 0.320 | 0.318 | 0.000 ± 0.342 | 4.13e<sup>-12</sup> |
| AccessFold | 0.317 | 0.507 | 0.390 | 0.397 | 0.349 ± 0.315 | 6.05e<sup>-06</sup> |
| PETcoFold* | 0.434 | 0.365 | 0.397 | 0.395 | 0.376 ± 0.352 | 9.24e<sup>-05</sup> |
| RNAfoldc | 0.384 | 0.582 | 0.463 | 0.469 | 0.446 ± 0.318 | 0.007 |
| NUPACK | 0.446 | 0.483 | 0.464 | 0.461 | 0.474 ± 0.355 | 0.001 |
| MXfold2c | 0.391 | 0.603 | 0.474 | 0.482 | 0.448 ± 0.320 | 0.004 |
| UFoldc | 0.422 | 0.545 | 0.476 | 0.476 | 0.455 ± 0.302 | 0.002 |
| bifold | 0.412 | 0.566 | 0.477 | 0.480 | 0.454 ± 0.325 | 0.014 |
| IntaRNA2.0 | 0.536 | 0.436 | 0.481 | 0.481 | 0.405 ± 0.377 | 9.75e<sup>-05</sup> |
| PairFold | 0.427 | 0.562 | 0.485 | 0.486 | 0.500 ± 0.339 | 0.007 |
| RNAmultifold | 0.423 | 0.582 | 0.490 | 0.493 | 0.494 ± 0.325 | 0.015 |
| RNAcoFold | 0.424 | 0.582 | 0.491 | 0.494 | 0.494 ± 0.325 | 0.015 |
| EternaFoldc | 0.473 | **0.618** | 0.512 | 0.519 | 0.498 ± 0.314 | 0.088 |
| SPOT-RNA2c* | 0.583 | 0.540 | 0.561 | 0.559 | 0.533 ± 0.331 | 0.012 |
| SPOT-RNAc | **0.620** | 0.551 | **0.583** | **0.583** | **0.582** ± 0.326 | |

Note: The overall F1-score is harmonic mean of precision and recall for all RNA pairs. PPV denotes positive predictive value. MCC denotes Matthews' correlation coefficient. The star \* denotes the use of evolution information. Methods with an ending of "c" indicate the use of chain concatenation for RNA-RNA interaction prediction. Median F1 means the median F1 value of single RNA. The P-value of a given method was computed by against the result from SPOT-RNAc.

As shown in Figure 1A, SPOT-RNA2c is the most accurate predictor at low sensitivity (<0.2). However, the overall PR curve given by SPOT-RNAc has the best performance. We can also measure the performance by the overall F1-Score for all RNA pairs and the median F1-Score for individual RNA pairs. The thresholds for determining F1-scores of SPOT-RNAc and SPOT-RNA2c in the test set were set according to the thresholds for producing the highest F1-scores in the validation dataset for SPOT-RNAc and SPOT-RNA2c, respectively. Table 1 confirmed the result from the PR curve that SPOT-RNAc achieved the best overall performance with an overall F1-Score of 0.583, outperforming SPOT-RNA2c (the overall F1-Score of 0.561) and EternaFold (F1-Score of 0.512). SPOT-RNAc improves over SPOT-RNA2c by more than 4% and outperforms other methods by over 14%, a pattern similarly observed in MCC values (Table 1).

Figure 1B presents the distribution of F1 scores for individual RNA pairs, including median, 25th, and 75th percentiles. SPOT-RNAc continues to achieve the best performance with the highest median F1 score of 0.582, outperforming the next best SPOT-RNA2c (median F1-score of 0.533) with a 10% improvement. The improvement of SPOT-RNAc over all methods are statistically significant except EternaFold (Table 1).

It’s important to assess how these methods perform on intra-molecular interactions, although not all RRI methods offer predictions for such interactions. We remove these RNAs without intra-base-pairing structures. This leads to 77 RNA chains. Figure 2A compares the PR curves or PR points given by 12 methods. Interestingly, PR curves indicate that SPOT-RNA2c now has the best performance. The overall performance according to overall F1-scores given by SPOT-RNA2c (Table 2) is the highest, surpassing the next best methods (SPOT-RNA2 without chain concatenation) by a 6% improvement in F1-score, and the third best (RNAmultifold or RNAcoFold) by 12%, with SPOT-RNAc ranking as the fifth best. This trend is consistent with MCC values.

**Figure 2:**
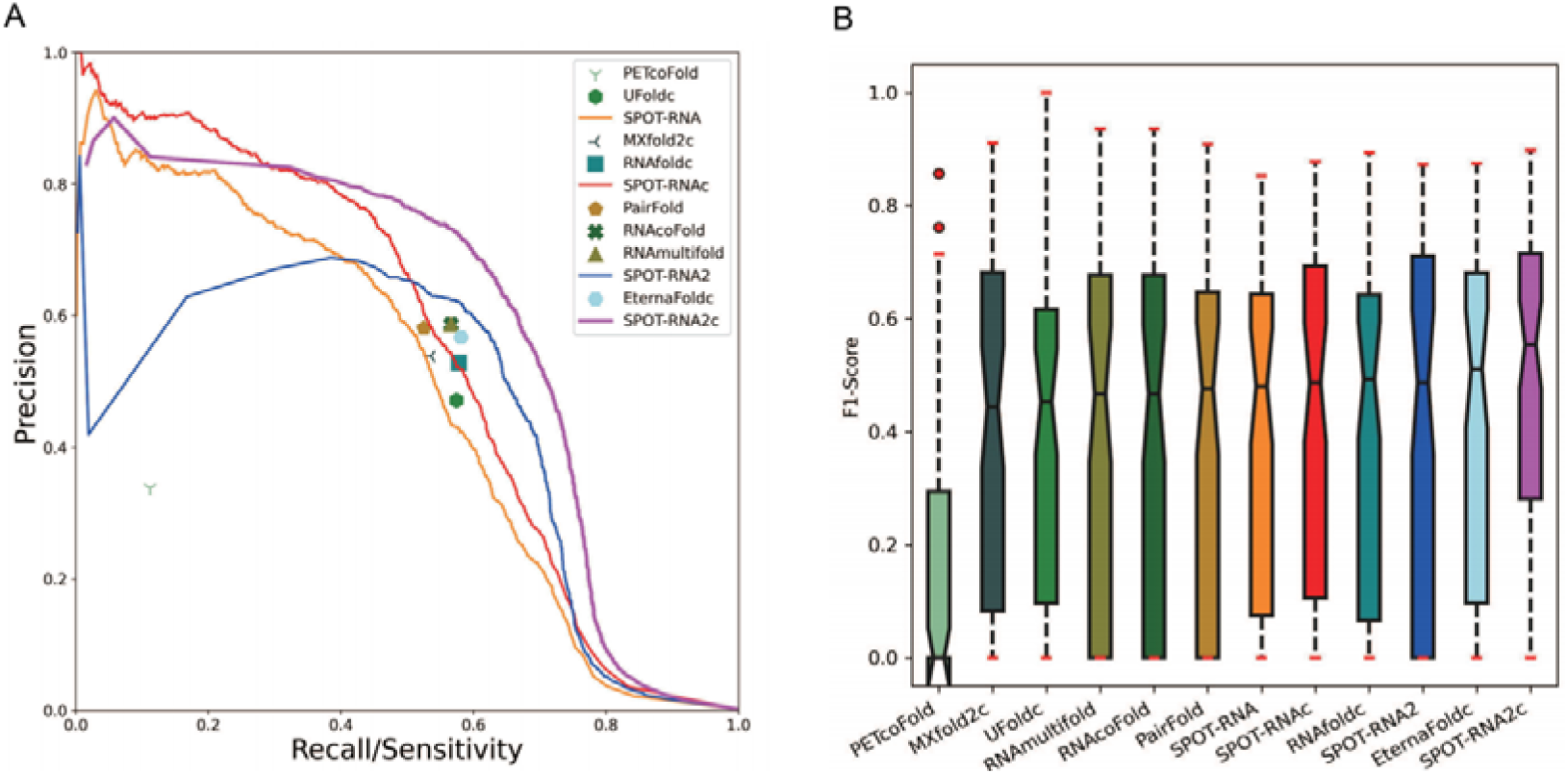
Performance Comparison of Intra-RNA Base Pair Prediction by 12 methods (A) Precision-recall curves and points illustrate the performance rankings. (B) A distribution of F1-scores for intra-RNA base pair prediction. SPOT-RNA and SPOT-RNAc represent intramolecular base pair prediction as single and concatenated chains, respectively. Evolution-based SPOT-RNA2c (or SPOT-RNA2) outperforms SPOT-RNAc (or SPOT-RNA) for intra-RNA base pairs.

**Table 2:** Performance Comparison of 12 Predictors for Predicting Intra-RNA Base Pairs on 77 unseen RNA structures that have intra secondary structure in 64 RNA-RNA Complex Structures.

| Methods | PPV/<br>Precision | Sensitivity<br>/Recall | Overall<br>F1-Score | MCC | Individual F1-Score Median ±<br>Std | P-value |
| --- | --- | --- | --- | --- | --- | --- |
| PETcoFold* | 0.338 | 0.112 | 0.168 | 0.194 | 0.000 ± 0.217 | 1.62e<sup>-15</sup> |
| UFoldc | 0.471 | 0.574 | 0.517 | 0.519 | 0.452 ± 0.278 | 5.33e<sup>-07</sup> |
| SPOT-RNA | 0.584 | 0.486 | 0.531 | 0.531 | 0.476 ± 0.271 | 1.51e<sup>-06</sup> |
| MXfold2c | 0.538 | 0.536 | 0.537 | 0.536 | 0.444 ± 0.291 | 0.0001 |
| RNAfoldc | 0.528 | 0.578 | 0.552 | 0.551 | 0.494 ± 0.287 | 1.06e<sup>-05</sup> |
| PairFold | 0.581 | 0.525 | 0.552 | 0.551 | 0.483 ± 0.308 | 8.15e<sup>-05</sup> |
| SPOT-RNAc | 0.653 | 0.493 | 0.562 | 0.567 | 0.481 ± 0.291 | 0.0005 |
| EternaFoldc | 0.567 | 0.581 | 0.574 | 0.573 | 0.511 ± 0.294 | 0.001 |
| RNAcoFold | 0.587 | 0.566 | 0.576 | 0.576 | 0.467 ± 0.305 | 0.001 |
| RNAmultifold | 0.587 | 0.566 | 0.576 | 0.576 | 0.467 ± 0.305 | 0.001 |
| SPOT-RNA2* | 0.589 | 0.629 | 0.608 | 0.607 | 0.489 ± 0.310 | 0.007 |
| SPOT-RNA2c* | 0.626 | 0.665 | 0.645 | 0.644 | 0.560 ± 0.294 | - |

Note: The overall F1-score is harmonic mean of precision and recall for all RNA pairs. PPV denotes positive predictive value. MCC denotes Matthew's correlation coefficient. The star \* indicates the use of evolution information. Std means standard deviation. Median F1 means the median F1
value of individual RNA chains. The P-value of a given method is calculated by against the result of SPOT-RNA2c.

Figure 2B further examines the distribution of F1-scores for individual RNAs. SPOT-RNA2c continues to lead with the highest median F1-score at 0.560, while EternaFold follows closely with the second-best median F1-score of 0.511. The differences in F1-score distributions between SPOT-RNA2c and other methods are all statistically significant, with a p-value of 0.001 when comparing SPOT-RNA2c to the second best EternaFold (Table 2).

Intuitively, a better intra-RNA base-pairing prediction should lead to a better inter-RNA base-pairing prediction. However, although SPOT-RNA2c has the best performance for intra-RNA interaction prediction, it is SPOT-RNAc with the best performance for inter-RNA interaction prediction. If we remove these RNA RRI pairs of which both chains do not have intra-base-pairing structures, this leads to 53 RRI pairs. Figure 3A compares intermolecular F1-scores for individual RNA pairs from SPOT-RNAc with the average intramolecular F1-scores. No correlation was found. Similar uncorrelated intra- and inter-RNA F1-scores are observed for SPOT-RNA2c (Figure 3B). This suggests that the evolution information contained in SPOT-RNA2c did contain the co-evolution information for predicting intra-RNA, but not inter-RNA interactions.

**Figure 3.**
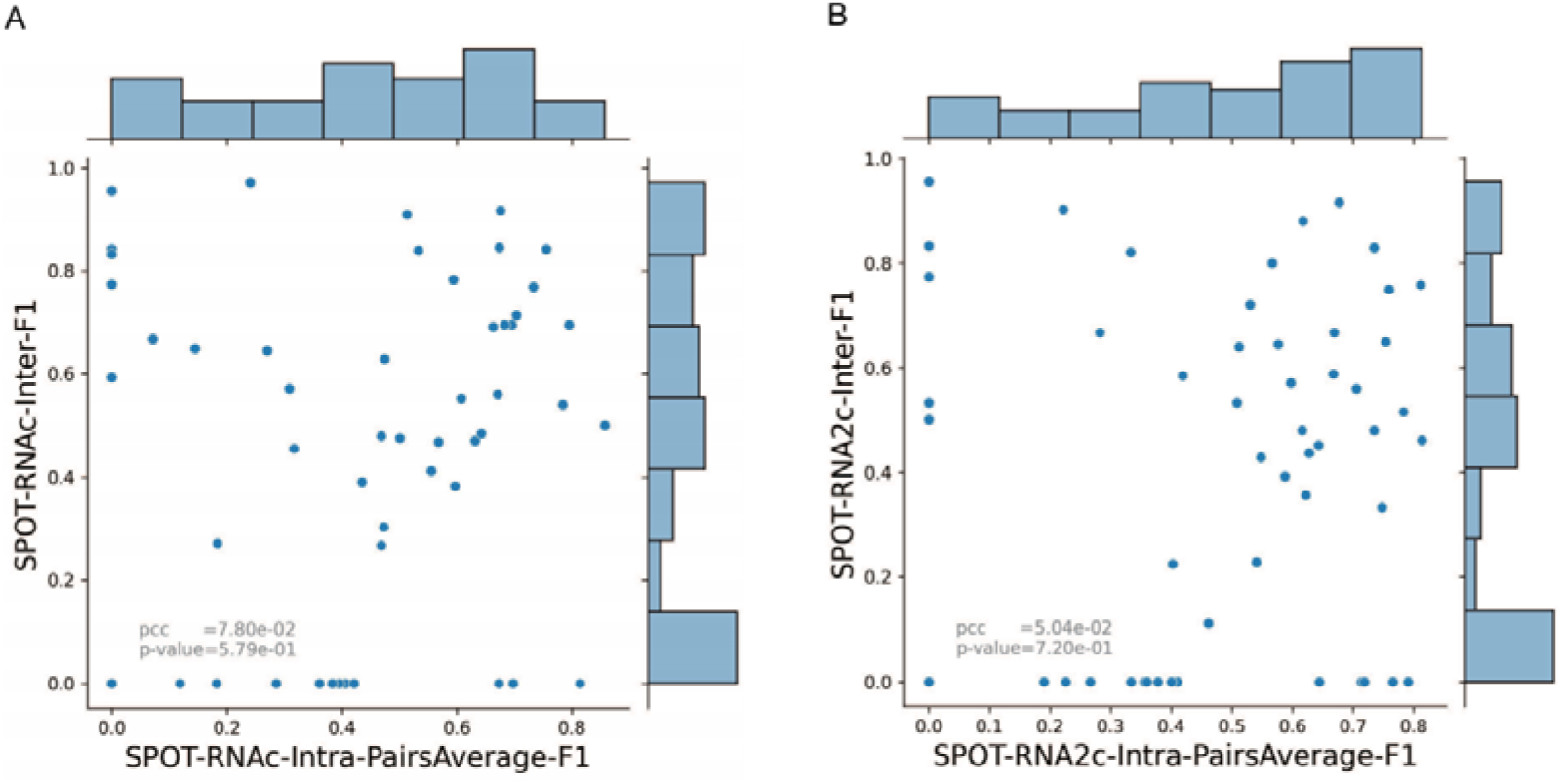
No Correlation between Inter-RNA F1-scores and Intra-RNA F1-scores. (A) Inter-RNA F1-scores versus the average Intra-RNA F1-scores of SPOT-RNAc for 53 RNA complex structures with intra-RNA base pairs for both chains. (B) Inter-RNA F1-scores versus Intra-RNA F1-scores of SPOT-RNA2c for 53 RNA complex structures.

We also assessed the impact of sequence concatenation on the intra-RNA base-pair prediction (i.e., SPOT-RNA versus SPOT-RNAc, SPOT-RNA2 versus SPOT-RNA2c). Table 2 shows that SPOT-RNAc/SPOT-RNA2c are better than SPOT-RNA/SPOT-RNA2 based on either overall F1-score or the median of individual F1-scores. In both cases, the difference is statistically significant with a p-value of 0.0007 between SPOT-RNA and SPOT-RNAc and a p-value of 0.007 between SPOT-RNA2 and SPOT-RNA2c. This indicates that knowing the binding partner improves the intra-RNA base pair prediction.

We observed that some inter-RNA (and intra-RNA) interactions were predicted with F1 score of 0 as shown in Figures 3A and 3B. To understand the reason behind these poor predictions, we examined F1-scores given by SPOT-RNAc as a function of sequence length (L1+L2) in Figure 4A. No obvious correlation was found. However, when we plotted F1-scores against the number of true inter-RNA base pairs divided by the square root of (L1*L2), a clear and strong correlation emerged with a Pearson’s correlation coefficient (PCC) of 0.464 (Figure 4B). Thus, poor predictions, including those with F1-scores of 0, can be attributed to the scarcity of inter-RNA contacts relative to the sequence lengths. This observation holds true for intra-RNA base pair prediction as well: intra-RNA interactions with F1-scores of 0 also involve very few intra-RNA base pairs (<5).

**Figure 4:**
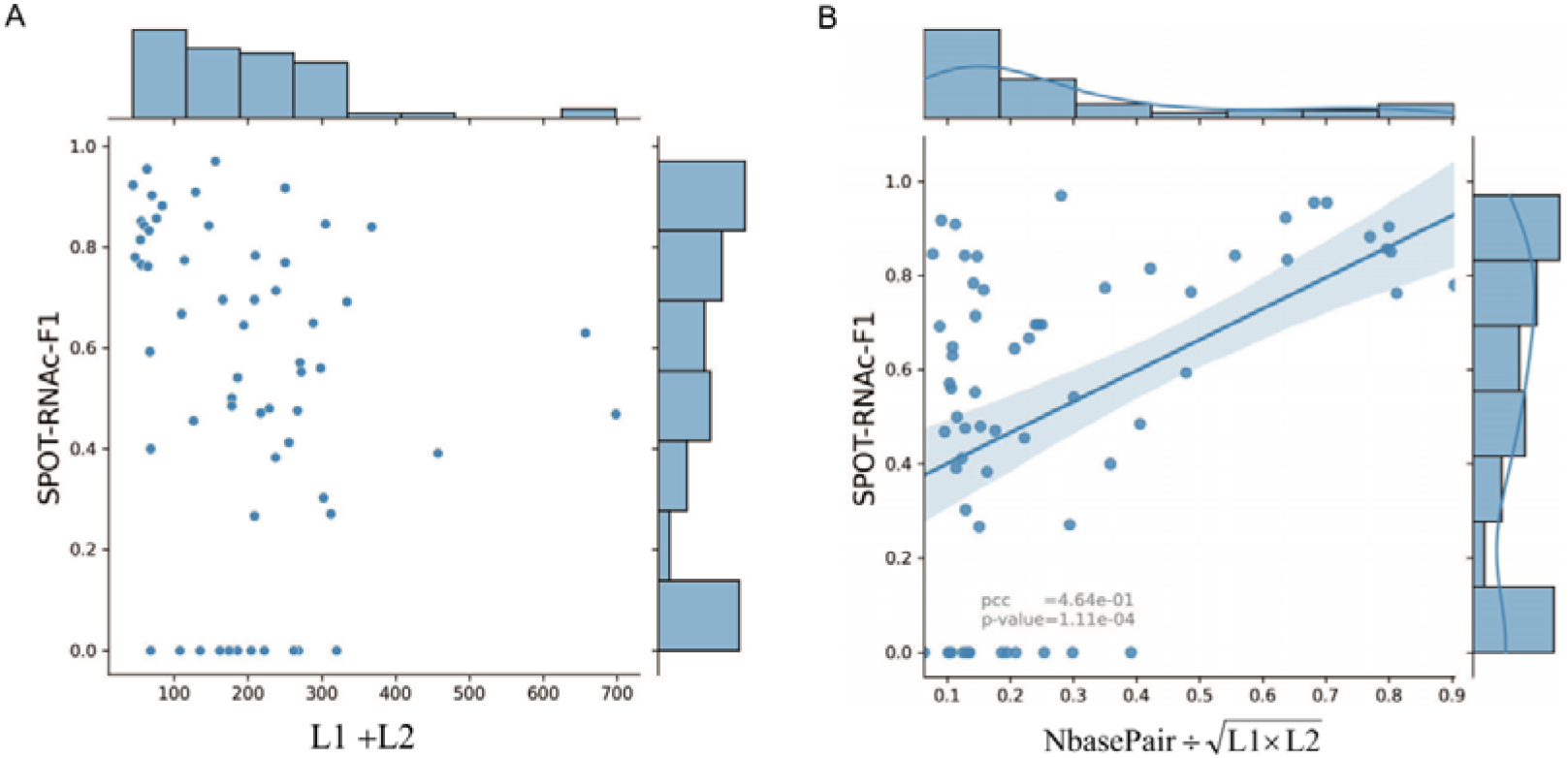
Relationship Between Inter-RNA Base Pairing Prediction and Sequence/Interaction Characteristics. (A) The inter-RNA F1 scores for individual RNA-RNA complexes from SPOT-RNAc plotted against the sum of sequence lengths (L1+L2). (B) The inter-RNA F1 scores for individual RNA-RNA complexes from SPOT-RNAc plotted against the normalized number of inter-RNA base pairs (the number of true inter-RNA base pairs divided by the 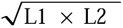). The performance does not correlate with sequence length but is related to the normalized number of inter-RNA base pairs.

One question is whether the method performance depends on RNA types. We classified 64 complexes into rRNA-containing complexes (only 8), snRNA/snoRNA/mRNA-containing complexes (40), and RNA complexes lacking intra-RNA interactions (16). SPOT-RNAc is consistently the best in each category as shown Supplementary Table S2 for the overall F1-score and MCC values. It is of note that even for RNA complexes lacking intra-RNA interactions (i.e. these extreme cases of unstructured RNAs are not employed for training in neither SPOT-RNA, nor SPOT-RNA2), SPOT-RNAc continues to provide the best overall F1-score and MCC values, although its performance for averaging over individual complexes is statistically similar to the those of some energy-based techniques. The consistency of SPOT-RNAc performance among different RNA types demonstrates that deep learning, even with a limited number of 3D structures for training, can yield generalizable models for inter-RNA base pair prediction.

Despite the attempt to remove structural homologs from training by structural alignment, some RNA families in training sets (rRNA and spliceosome complexes) were still included in the above 64-complex test sets because RNAs in the same family do not necessarily have an identical structure. We further examined the effect of excluding-training families in intra-base pair prediction by establishing new datasets. They were obtained by removing training/validation families from 64 test complexes and locating additional new RNA complexes (up to 2024/02/28) (See methods). These two datasets are mixed and separated into 15 hetero dimeric complexes and 13 homo-dimeric complexes. For 15 hetero-dimer complexes, although the difference between SPOT-RNAc and some methods are statistically insignificant, SPOT-RNAc remains the best performance among all the methods compared for the overall performance (Table 3), whereas SPOT-RNA2 remains the best for intra-RNA base-pair prediction (Supplementary Table S3) with an overall F1-score of 0.589, followed by RNAmultifold (0.586) and RNAcoFold (0.586). If ranked by median F1-score, SPOT-RNA2c remains the best for intra-RNA interaction predition. Most did poorly with a median F1-score of 0 except SPOT-RNA2c, due to low number of intra-RNA contacts for this set of the data. All also did very poorly with median F1-score of 0 for inter-RNA base-pair prediction in the 13-homodimer set (Supplementary Table S4), whereas SPOT-RNA2 is the best performance with an overall F1-score of 0.750, followed by SPOT-RNA (0.587) and RNA-RNA2c (0.529) for intra-RNA base-pair prediction (Supplementary Table S5).

**Table 3.**
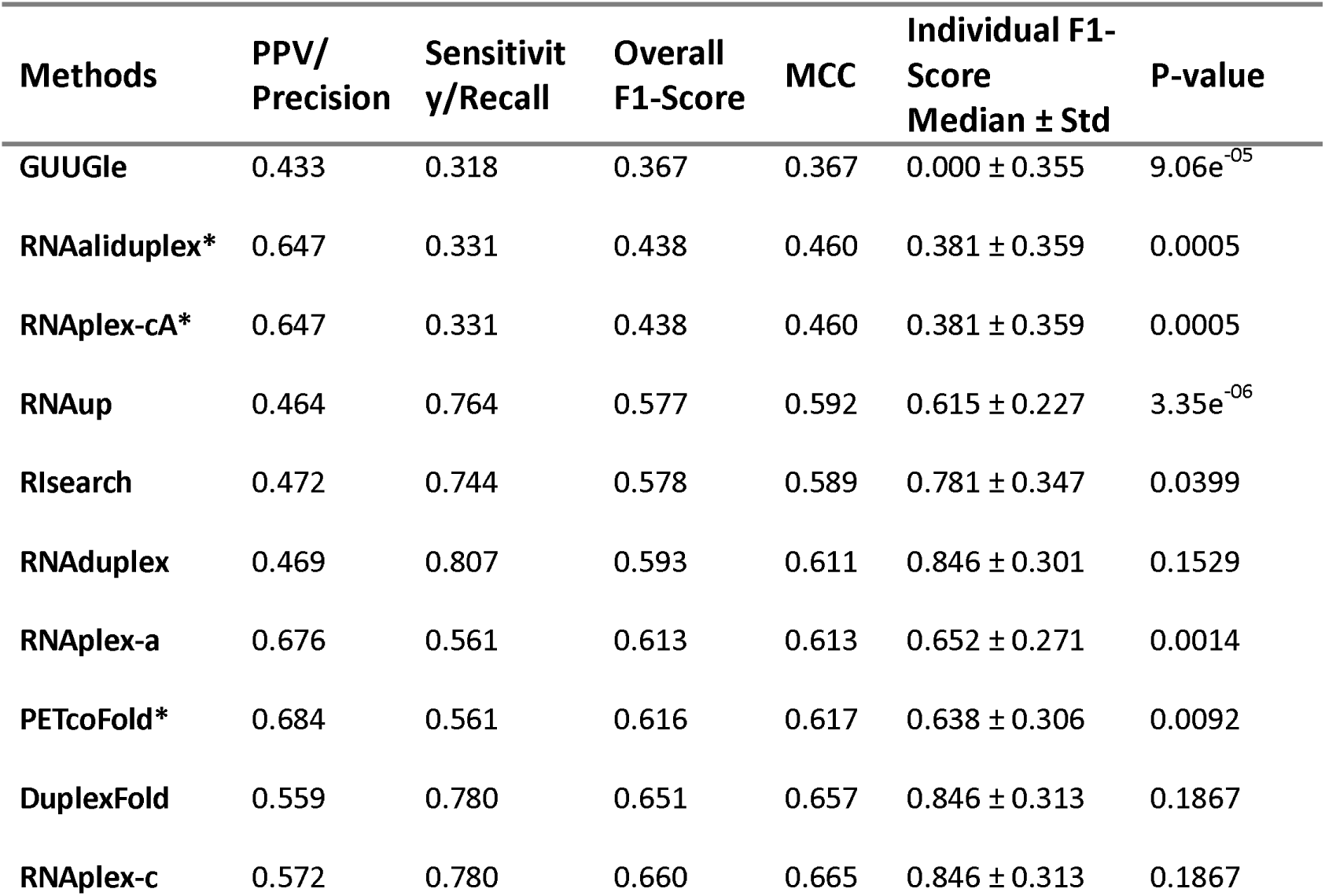
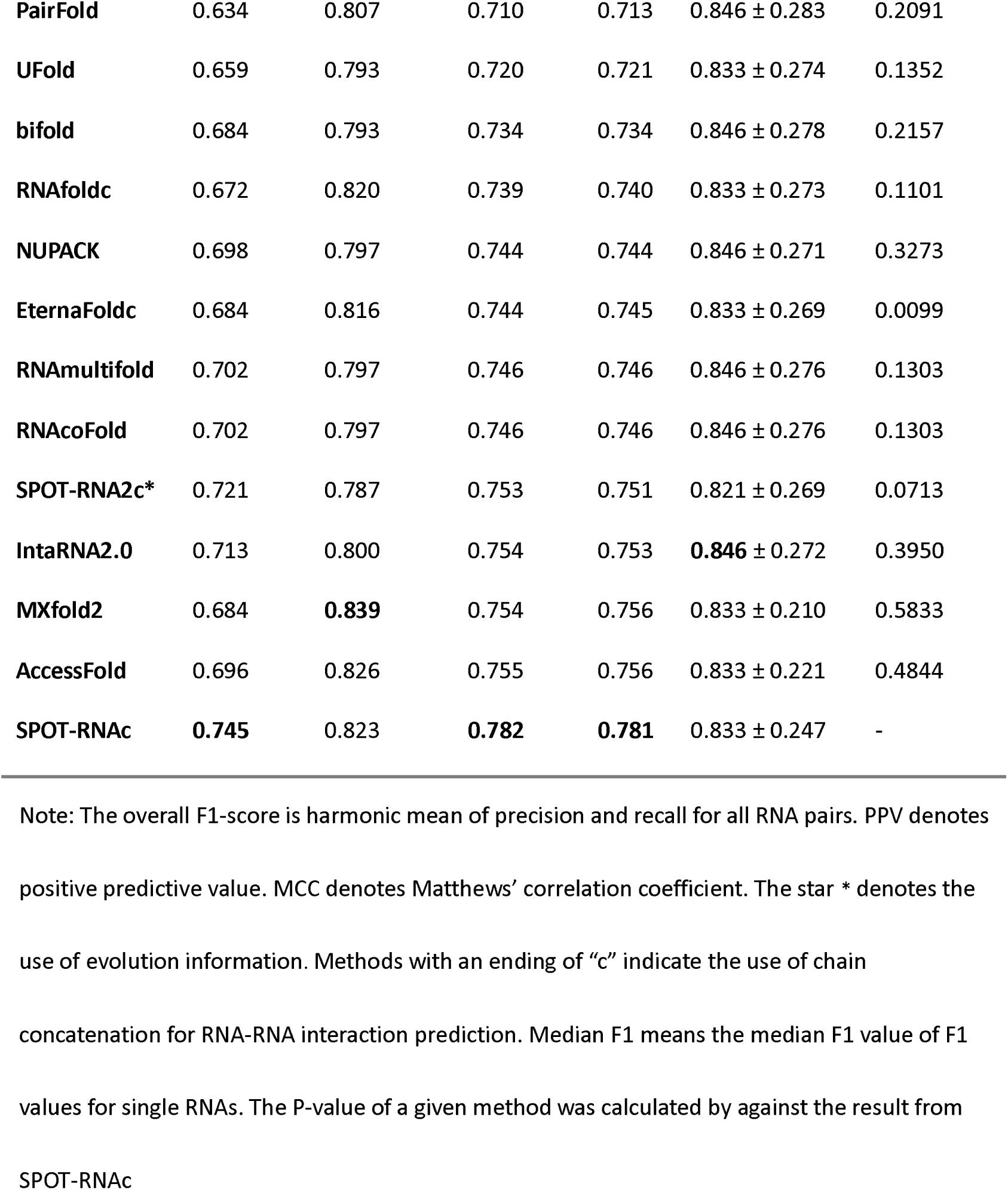
Performance comparison of all predictors on 15 hetero-dimer complexes of inter-RNA base pairs that are not in RNA families containing in TR0/VL0 (bpRNA) and TR1/VL1 for training SPOT-RNA and SPOT-RNA2.

| Methods | PPV/<br>Precision | Sensitivity/<br>Recall | Overall<br>F1-Score | MCC | Individual F1-Score<br>Median ± Std | P-value |
| --- | --- | --- | --- | --- | --- | --- |
| GUUGle | 0.433 | 0.318 | 0.367 | 0.367 | 0.000 ± 0.355 | 9.06e<sup>-05</sup> |
| RNAaliduplex* | 0.647 | 0.331 | 0.438 | 0.460 | 0.381 ± 0.359 | 0.0005 |
| RNAplex-cA* | 0.647 | 0.331 | 0.438 | 0.460 | 0.381 ± 0.359 | 0.0005 |
| RNAup | 0.464 | 0.764 | 0.577 | 0.592 | 0.615 ± 0.227 | 3.35e<sup>-06</sup> |
| Rlsearch | 0.472 | 0.744 | 0.578 | 0.589 | 0.781 ± 0.347 | 0.0399 |
| RNA duplex | 0.469 | 0.807 | 0.593 | 0.611 | 0.846 ± 0.301 | 0.1529 |
| RNAplex-a | 0.676 | 0.561 | 0.613 | 0.613 | 0.652 ± 0.271 | 0.0014 |
| PETcoFold* | 0.684 | 0.561 | 0.616 | 0.617 | 0.638 ± 0.306 | 0.0092 |
| DuplexFold | 0.559 | 0.780 | 0.651 | 0.657 | 0.846 ± 0.313 | 0.1867 |
| RNAplex-c | 0.572 | 0.780 | 0.660 | 0.665 | 0.846 ± 0.313 | 0.1867 |
| **PairFold** | 0.634 | 0.807 | 0.710 | 0.713 | 0.846 ± 0.283 | 0.2091 |
| **UFold** | 0.659 | 0.793 | 0.720 | 0.721 | 0.833 ± 0.274 | 0.1352 |
| **bifold** | 0.684 | 0.793 | 0.734 | 0.734 | 0.846 ± 0.278 | 0.2157 |
| **RNAfoldc** | 0.672 | 0.820 | 0.739 | 0.740 | 0.833 ± 0.273 | 0.1101 |
| **NUPACK** | 0.698 | 0.797 | 0.744 | 0.744 | 0.846 ± 0.271 | 0.3273 |
| **EternaFoldc** | 0.684 | 0.816 | 0.744 | 0.745 | 0.833 ± 0.269 | 0.0099 |
| **RNAmultifold** | 0.702 | 0.797 | 0.746 | 0.746 | 0.846 ± 0.276 | 0.1303 |
| **RNAcoFold** | 0.702 | 0.797 | 0.746 | 0.746 | 0.846 ± 0.276 | 0.1303 |
| **SPOT-RNA2c\*** | 0.721 | 0.787 | 0.753 | 0.751 | 0.821 ± 0.269 | 0.0713 |
| **IntaRNA2.0** | 0.713 | 0.800 | 0.754 | 0.753 | **0.846** ± 0.272 | 0.3950 |
| **MXfold2** | 0.684 | **0.839** | 0.754 | 0.756 | 0.833 ± 0.210 | 0.5833 |
| **AccessFold** | 0.696 | 0.826 | 0.755 | 0.756 | 0.833 ± 0.221 | 0.4844 |
| **SPOT-RNAc** | **0.745** | 0.823 | **0.782** | **0.781** | 0.833 ± 0.247 | - |

Note: The overall F1-score is harmonic mean of precision and recall for all RNA pairs. PPV denotes positive predictive value. MCC denotes Matthews' correlation coefficient. The star \* denotes the use of evolution information. Methods with an ending of "c" indicate the use of chain concatenation for RNA-RNA interaction prediction. Median F1 means the median F1 value of F1 values for single RNAs. The P-value of a given method was calculated by against the result from SPOT-RNAc

We selected one example to illustrate SPOT-RNA’s performance in RRI prediction. Figure 4A displays predicted and actual base-pairing maps for the subunits L2a rRNA complexed with L3b rRNA from Chlamydomonas reinhardtii mitoribosome (PDB ID 7PKT, chain ID 2 and chain ID 3). Figure 5A displays the intra- and inter-base-pairing maps of these two RNAs with Figure 5B for inter-base-pairing maps only. The F1 scores for intra-RNA base pairs are 0.32 for L2a rRNA and 0.30 for L3b rRNA, while the F1-score for inter-RNA base pairs is 0.571. Correctly predicted base pairs are highlighted with red dots in Figures 5B and 5C, and their 3D locations were shown as red-colored bases in Figure 5C. For this complex structure, precision for inter-RNA base pairs is 0.435, and sensitivity is relatively high (0.769).

**Figure 5:**
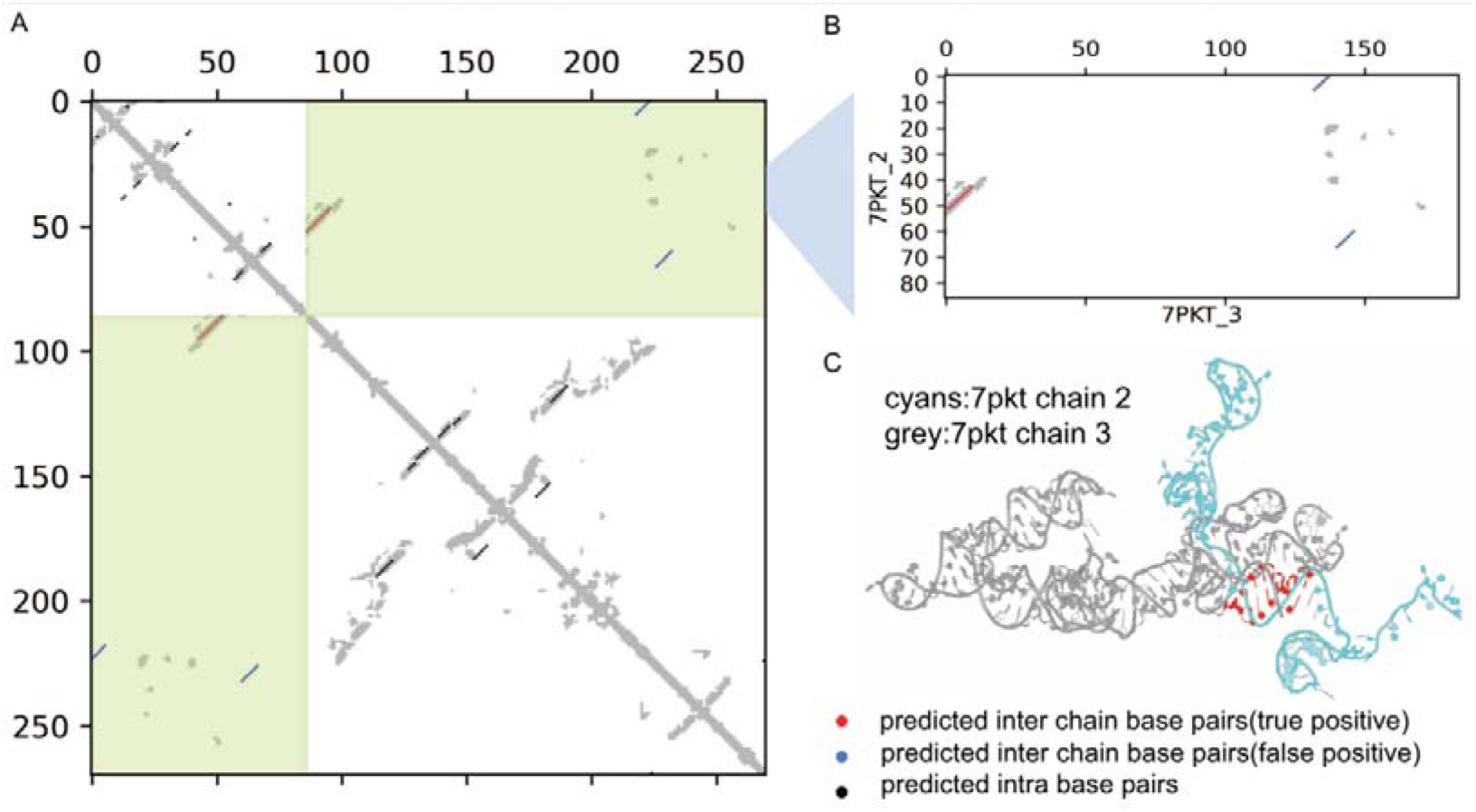
Accurate Prediction of Key Inter-RNA Base-pairing Contacts in RNA Complexes by SPOT-RNAc. Complex structure of large subunits of the Chlamydomonas reinhardtii mitoribosome (L2aRNA and L3bRNA in PDB ID 7PKT) with true distance-contact map and predicted intra and inter-RNA base pairs (A), inter-RNA base pairs only (B), and 3-D structure (C). Predicted intra-base pairs and inter-base pairs are denoted by black and red dots, respectively, in the base-pairing maps. In the 3-D structure (C), correctly predicted inter-RNA base pairs are highlighted in red.

## Discussion

This study represents a comprehensive benchmark for assessing more than 20 methods in predicting RNA-RNA interactions. Previous benchmarks were constrained to interactions involving small RNAs without intra-RNA base pairs. For instance, Lai and Meyer ^13^ compared 14 RRI methods based on experimentally confirmed interactions in fungal snoRNA-rRNA and bacterial sRNA-mRNA pairs. Umu and Gardner ^12^ examined 15 RRI methods using a dataset focused on short linear base-pair matching. Antonov et al.^38^ compared 13 RRI methods on mammalian lncRNAs with experimentally proven hybridizations. In contrast, our study presents the first comprehensive benchmark of 23 RRI prediction methods using known RRI interactions derived from 3D structures at the base pair level. Notably, most of these complexes (53/64) include RNAs with 3D structures and intra-RNA base pairs. Additionally, this study marks the first inclusion of deep-learning based methods for comparisons, utilizing chain concatenation.

In contrast to proteins, the available non-redundant data for RNA structures is limited. This raises concerns about the generalizability of deep learning models trained on such limited data. Previous studies by Szikszai et al ^39^ and Qiu ^40^ highlighted the challenges of deep-learning models when applied to unseen families not present in the training and validation sets. SPOT-RNA and SPOT-RNA2 were initially trained with a set of bpRNA dataset after removing all sequences in bpRNA with known 3D structures by CD-HIT (80% sequence identity cut-off), followed by transfer learning with 3D-structure-derived base pairs. To evaluate the adaptability of SPOT-RNA and SPOT-RNA2 beyond their training and validation data, we placed them to the test of untrained inter-RNA interactions. To avoid possible “over-trained” intra-RNA interactions affecting prediction of inter-RNA interactions, we excluded all single-chain structures in the complex test set with the structural similarity score (TM-score)≥0.3, compared to those in the training and validation sets of SPOT-RNA. Interestingly, the performance of intra-RNA base-pair prediction does not correlate with the performance of inter-RNA base-pair prediction (Figure 3A). Lacking correlation suggests that the performance for predicting inter-RNA interactions is independent of the performance for predicting intra-RNA interactions and the dataset for inter-RNA interactions can be employed as an independent test set for the methods trained for predicting intra-RNA interactions. Despite the above precaution, training/validation families were not entirely excluded because RNAs in the same family may not have an identical structure. Thus, we further established two family-excluded test sets (15 hetero-dimeric RNAs and 13 homo-dimeric RNAs). SPOT-RNA and SPOT-RNA2 remain best performance for inter and intra-RNA base-pair prediction, respectively, except for inter-base-pair prediction for homodimeric RNAs, for which no methods did well. The generalization capability of SPOT-RNA is further illustrated with its robust performance across different RNA types, even for RNAs with few intra-RNA structures (Supplementary Table S2).

SPOT-RNA2, which incorporates evolution and co-evolution information, outperforms SPOT-RNA for intra-RNA base pairs, aligning with previous findings (Table 2) ^35^. However, SPOT-RNA2c underperforms SPOT-RNAc for inter-RNA base pairs. Notably, SPOT-RNA2 exhibits a positive correlation with the number of effective homologous sequences for intra-RNA base-pair prediction (PCC=0.5, p-value=2×10^-6^, Supplementary Figure 1C), but this correlation nearly diminishes for SPOT-RNA2c for intra-RNA base pairs (PCC=0.27, p-value=0.05) and turned negative for inter-RNA base pair prediction (PCC=-0.3, p-value=0.02). This suggests that using linked sequences in SPOT-RNA2c for homology search may have provided harmful information for inferring inter-RNA interactions. In the future, it may be necessary to utilize sequences from the same species for homology searches, as co-evolution information can only be detected through inter-species comparisons via Multiple Sequence Alignment (MSA) pairing as has been done for proteins^41^. However, the failure of SPOT-RNA2 in predicting inter-RNA interactions between homodimers indicates that the coevolution signals between RNAs may be too weak to capture.

For predicting RNA-RNA interactions, we concatenated two chains (A and B) as a single chain. To eliminate artificial sequence-order dependence, we predicted results for both AB and BA chains and then calculated the average. Interestingly, we found that in some cases, one sequence order (e.g., BA) outperformed the other (i.e., AB). Upon closer examination, we discovered for some RNA pairs that the order with a shorter separation in sequence positions for contacting base pairs tended to perform better. This observation is intuitive, as longer-range interactions are inherently more challenging to predict. However, there were some outliers deserving further studies.

## Methods

### Benchmark datasets

We retrieved all RNA structures from the Protein Data Bank (PDB) in March 2023^10^ and specifically selected structures featuring two RNA chains with a minimum of 5 inter-RNA base pairs. The identification of base pairs was carried out using DSSR^42^. For the base-pair prediction in complex structures, interacting base pairs inside the complex are positive samples, and unpaired bases between them are naturally classified as negative samples. To eliminate redundancy, we applied CD-HIT-EST^43^, removing binding pairs with over 80% sequence identity between either chain. This initial step resulted in 155 unique RNA-RNA interaction (RRI) pairs.

To ensure stringency, we further filtered out RRI pairs that exhibited single-chain structural similarities with any RNAs in the SPOT-RNA training set, defined by TM-score ≥0.3 using RNA-align^44^ with the length of the query sequence for normalization. This rigorous process yielded a final benchmark set comprising 64 unique RRI pairs. The PDB IDs of the benchmark set can be found in Supplementary Table S6. To further build a dataset excluding RNA families employed in training/validating SPOT-RNA, we identified 14 complexes out of 64 test complexes whose RNAs do not belong to any RFAM families with RNAs contained in training/validation TR0/VL0 (bpRNA) and TR1/VL1 (RNAs with 3D structures) sets of SPOT-RNA.

To expand this dataset, we re-downloaded all RNA complexes on 2024/02/28 from the protein data bank and processed by DSSR. Potential complexes were extracted from asymmetric units and biological assemblies and then filtered such that only structures with 6 or more intermolecular and 3 or more intramolecular base pairs (canonical and non-canonical) per subunit were retained in the dataset. Any complex with a subunit of length >400 nucleotides (almost exclusively rRNA and snRNA) or containing only subunits with length less than 25 nucleotides were removed from the dataset. Remaining RNA were filtered against the SPOT-RNA training/validation sets (TR0/VL0/TR1/VL1) using blastn (-task blastn-short) with an e-value cutoff of 0.001 and using RNA-align with a cutoff of 0.45. Identical duplicates were removed by selecting structures with the largest number of base-pairing interactions. Two RNA (7r6q – rRNA and 6ltp - crRNA) were also removed after manual inspection. Finally, we found that except for three of the new complexes, all other complexes are homooligomers and among the original complexes, only two of them are homooligomeric. Therefore, we integrated these two datasets, resulting in a homo-dimer data set containing 13 complexes and a hetero-dimer data set containing 15 complexes.

### Performance evaluation

We evaluated performance using common metrics: Recall (sensitivity), Precision, and the F1 score. PPV/Precision is TP/(TP+FP), Sensitivity/Recall is TP/(TP+FN), and the F1 score is 2(Recall*Precision)/(Recall + Precision). Here, TP, FP, and FN are true positive, false positive and false negative, respectively. We also calculated Matthew’s correlation coefficient (MCC) to provide a balanced measure as below:

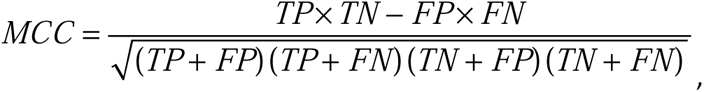

where TN denotes true negatives. MCC considers true negatives (TN) and measures the correlation between expected and observed classes. It ranges from 0 (no correlation) to 1 (highest correlation).

### Methods for interaction prediction

We summarized the methods compared in Table 4, with detailed settings for each method provided in the Supplementary method description in the Supplementary Material. As per Lai and Meyer^13^, we categorized the algorithms into four types: 1) ‘Interaction only’ methods predict intermolecular hybridization, ignoring RNA secondary structures; 2) ‘Accessibility’ methods consider RNA secondary structures using a partition function for unpaired probability; 3) ‘Concatenation’ algorithms treat two input sequences as a single chain and predict a joint secondary structure; and 4) ‘Complex joint’ methods also predict joint secondary structures but without concatenating the input sequences.

**Table 4.**
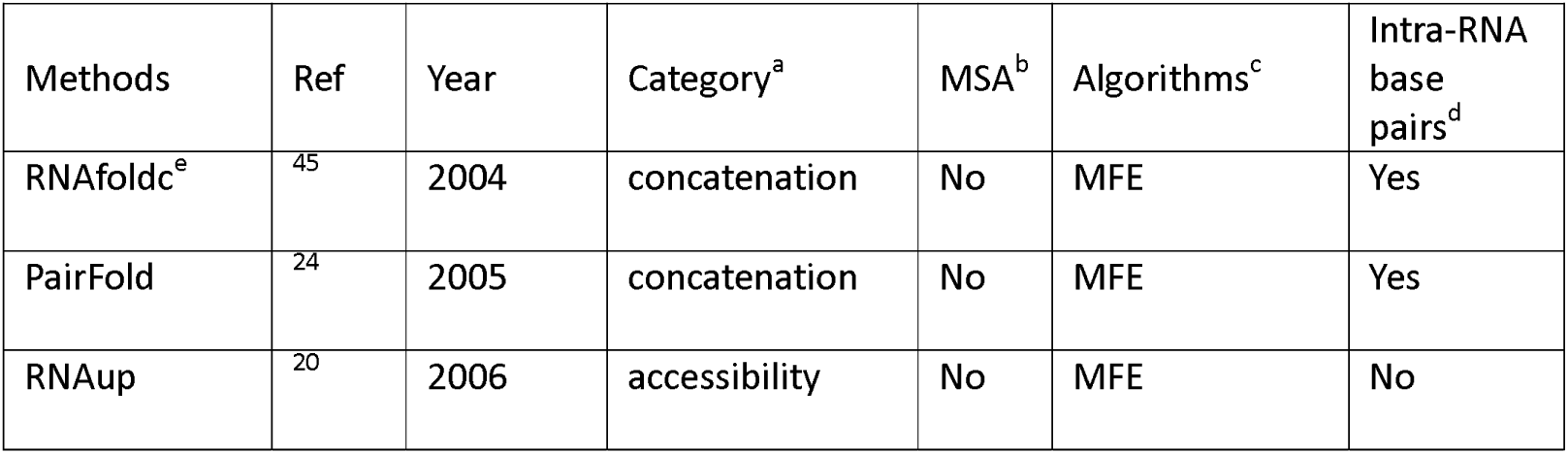
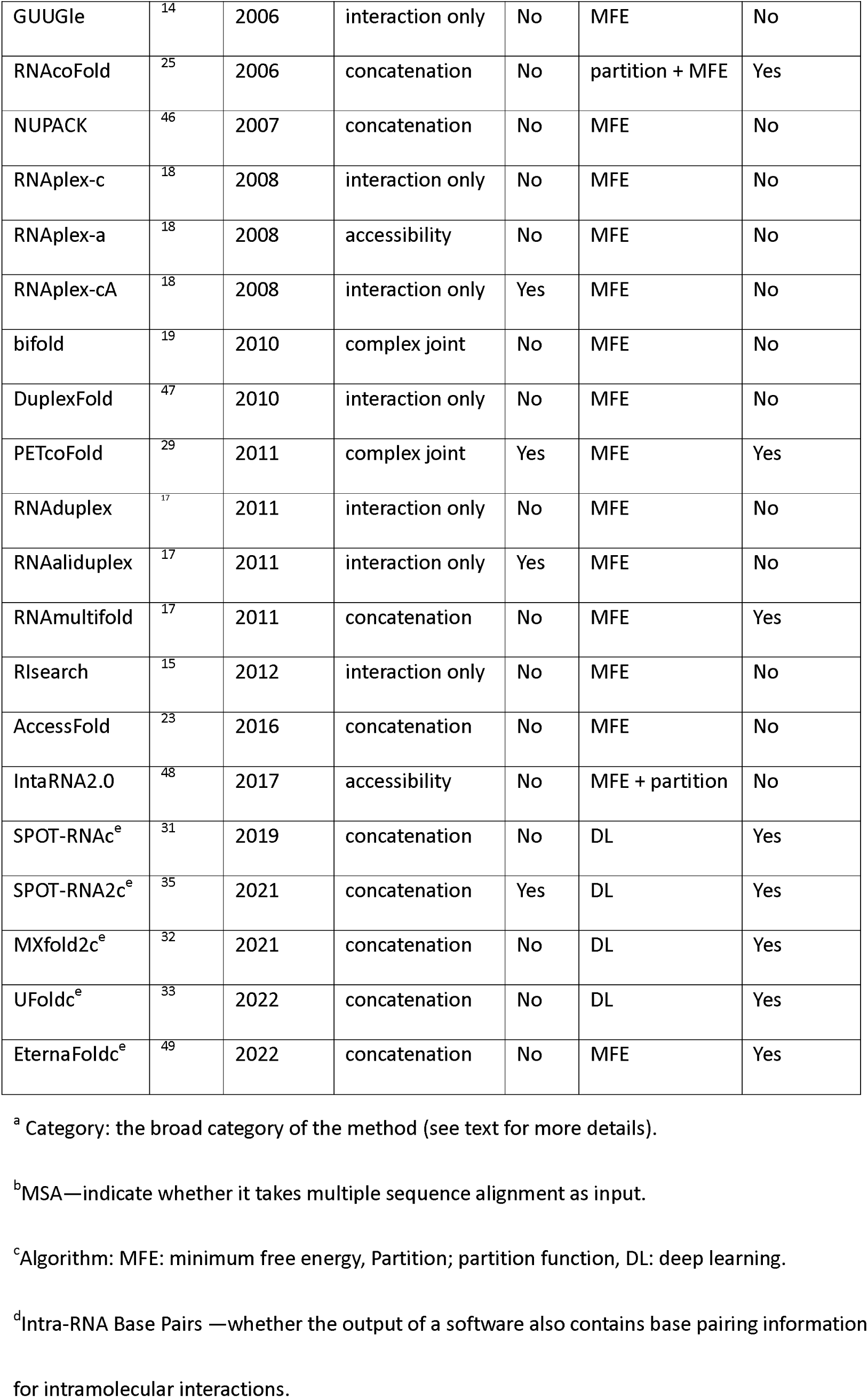
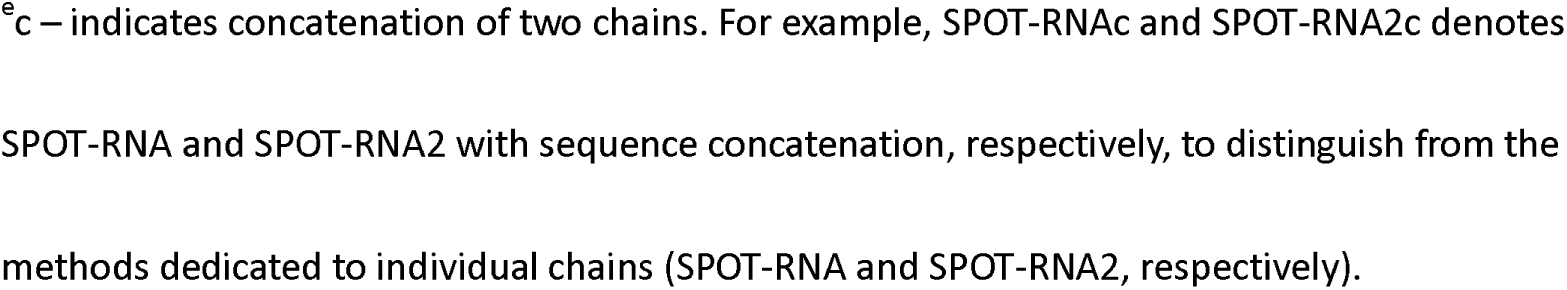
RRI interaction tools employed in this study for comparison, listed according to the year of publication along with their categories, the use of evolution information (MSA), the algorithm, and the capability of predicting intra-RNA base pairs.

We used concatenation for comparing recent deep-learning methods with traditional free-energy-based methods. When dealing with concatenated chains, we made predictions for both sequence orders (AB and BA) and reported the average if probabilities were predicted. For those methods providing a two-state prediction, we considered the union of base pairs predicted for both sequence orders as a positive prediction (i.e. a positive prediction from either AB or BA concatenation will be considered as positive).

We experimented with and without a three-nucleotide linker (AAA, UUU, CCC, or GGG) and found that direct concatenation without any linker yielded slightly better results, although not statistically significant compared to AAA/CCC linkers (Supplementary Table 1). Therefore, we report results based on direct concatenation without any linkers.

### Homology search

For these methods that do not predicted RRI by employing a linked chain such as RNAplex-cA, PETcoFold, RNAaliduplex and SPOT-RNA2, we searched the homologs using the single chain by RNAcmap3^50^ which is based on MARS database (a comprehensive database by including the RNAcentral, MG-RAST, GWH, MGnify and NCBI’s nucleotide databases). For SPOT-RNA2c using a link chain, we searched the homologs with the linked chain by RNAcmap3. The median number of homologous sequences found is 1467 for single chains and 940 for linked chains.

## Data and code availability

All data and codes are available at https://github.com/meilanglang/RNA-RNA-Interaction

## Acknowledgement

We express our gratitude to Dr. Jaswinder Singh for valuable discussions and insights. This research received support from the National Natural Science Foundation of China (Grant # 22350710182), the Shenzhen Science and Technology Program (Grant No. KQTD20170330155106581) and benefited from access to the supercomputing resources at Shenzhen Bay Laboratory.

